# Correlation Detection with and without the Theories of Conditionals: A model update of Hattori & Oaksford (2007)

**DOI:** 10.1101/247742

**Authors:** Tatsuji Takahashi, Kuratomo Oyo, Akihiro Tamatsukuri, Kohki Higuchi

## Abstract

We view observational causal induction as a statistical independence test under rarity assumption. This paper complements the two-stage theory of causal induction proposed by Hattori and Oaksford (2007) with a computational analysis. We show that their dual-factor heuristic (DFH) model has a rational account as the square root of the index of (non-)independence under extreme rarity assumption, contrary to the criticism that the DFH model is non-normative (e.g., Lu et al., 2008). We introduce a model that considers the proportion of assumed-to-be rare instances (pARIs), which is the probability of biconditionals (according to several theories of compound conditionals) and can be seen as a simplified version of the DFH model. While being a single conditional probability, pARIs approximates the non-independence measure, the square of DFH. In reproducing the meta-analysis in Hattori and Oaksford (2007), we confirm that pARIs and DFH have the same level of descriptive adequacy, and that the two models have the highest fit among more than 40 models. Then, we critically examine the computer simulations which were central to the rational analysis in Hattori and Oaksford (2007). We point out two problems in their simluations: samples in some of the simulations being restricted to generative ones, and indefinite values of models because of the small samples. In the light of especially the latter problem of definability, pARIs shows higher applicability.

## 1. Introduction

Causality is with no doubt one of the bases of human cognition (see e.g., Sloman and Lagnado (2015); Lake, Ullman, Tenenbaum and Gershman (2017)). Notwithstanding the elusive nature of causal relationships that cannot be directly observed but rather constructed through inductive inference by internal observers under uncertainty (e.g., Gunji, Takahashi and Aono (2004)), animals (including humans) have been remarkably successful in adapting to new environments through predictions of the future, manipulation of the environment, diagnosis of causes, and explanation of the reasons for or attribution of rewards, all based on learning and inference of causal relationship (Griffiths and Tenenbaum, 2009).

Efficient dealing with causal relationships has a high importance as for development of intelligent, learnable and autonomous systems (Lake et al., 2017). As the scope of the machine intelligence becomes broader, enabling the robots and agents to act in parallel with humans, the problems facing them become human-like and human-level as well. Although there is no necessity for agents to solve the problems just as humans do. There will be more and more problems facing the agents that are human-like and also similar in small amounts of input, instead of big data, where agents need to learn from small pieces of information given in real time. As humans’ inferences actually often work well, it is worth learning from it (Griffiths and Tenenbaum, 2009). Especially considering that the performance of human reasoning heavily depends on the existence of “mental mechanisms” that are causal models of the task to solve (Sloman, 2005; Chater and Oaksford, 2006), causal inference has high importance in human-like AI research as well (Lake et al., 2017).

In this study, inheriting the theoretical framework for causal induction by Hattori and Oaksford (2007), we critically examinine their model (DFH: dual-factor heuristics) and the simulations, and propose a simpler model (pARIs: proportion of assumed-to-be rare instances) that solves the problems that were implicit in their study (Takahashi, Kohno and Oyo, 2010). DFH and pARIs have similar biconditional forms; the difference consists in whether the biconditional form was derived from the limiting operation from the normative correlation coefficient (DFH) or derived from theories of conditionals and compound conditionals (pARIs), as first suggested by David Over in a JSPS-ANR CHORUS Franco-Japanese project (BTAFDOC) meeting on 5th March 2012 in Paris.

## 2. Causal induction

There are at least two kinds of causal inference: inductive formation of causal relationships from observation of co-occurrence of events, and testing the supposed relationships through experimentation/intervention, The first, most inductive kind is the focus of this study.

### 2.1. The two stage theory of causal induction: Observation then intervention

We adopt the two stage theory of causal induction (Hattori and Oaksford, 2007; Hattori, Over, Hattori, Takahashi and Baratgin, 2016; Hattori, Hattori, Over, Takahashi and Baratgin, 2017) as the theoretical framework in this study. According to the theory, in humans’ causal induction in realstic environments abound with innumerable candidate causes and effects, and people first observe co-occurrences between events to determine the most relevant-seeming cause-effect pairs. From pure observation, especially only with co-occurrence, it is usually impossible to distinguish between correlation and causation. However, in many cases, causally related events are correlated. Hence, detecting correlation and filtering out irrelevant event pairs can be considered the goal of the first, observational stage of causal inference. The interventional stage comes next, in which experimentation can be used to test whether the distilled pairs are causal or spurious.

Because of the abundance of events, together with limits to mental computation, spatio-temporal observation scope and working memory, it is rational to use some simplified strategy in the first stage to detect correlated pairs of events that would be relatively rare among numerous irrelevant pairs. It is in this sense that the first, observational stage is called heuristic, whereas the second, interventional stage is called analytic. The most well-organised example of the second, interventional stage is modern scientific experiments. The two stages ordered this way is expected to reduce the cost of intervention and experimentation. The distinction between the two stages, especially in terms of modelling, has gradually been established, both theoretically (Hattori et al., 2016) and empirically (Hattori et al., 2017).

### 2.2. Models of causal induction with one cause and one effect

The experimental and modelling framework of causal induction for this study is that agents form the causal strength from *C* to *E* based on the four frequencies in Table 1. *E* is the effect in focus and *C* is a candidate cause of *E*. We also assume that the models have the type of function with the domain of four natural numbers—*N*(*C, E*), *N*(*C*, ¬*E*), *N*(¬*C, E*) and *N*(¬*C*, ¬*E*)—as in Table 1, to the co-domain of an interval of real numbers: *f* : 𝒩^4^ → *V*. *V* is usually the unit interval [0, 1] for probability or [−1, 1] for indices like correlation coefficients. In the observational stage, the basic index on this table is the four-fold correlation coefficient *ϕ*:

**Table 1:** The 2×2 contingency table for elemental causal induction. Ω denotes the universe.

| | Effect ($E$) | No effect ($\neg E$) | Marginal frequency |
| --- | --- | --- | --- |
| Cause ($C$) | $N(C, E)$ | $N(C, \neg E)$ | $N(C)$ |
| No cause ($\neg C$) | $N(\neg C, E)$ | $N(\neg C, \neg E)$ | $N(\neg C)$ |
| Marginal frequency | $N(E)$ | $N(\neg E)$ | $N(\Omega)$ |

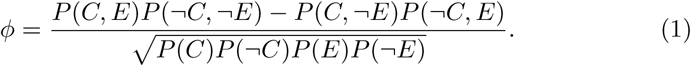

The most representative model of causal induction is Δ*P*(Jenkins and Ward, 1965):

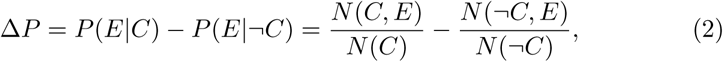

which is the increase in the probability of *E* caused by the presence of *C* relative to *C*’s absence. This model formalises the basic idea of scientific experiments. For example, the first term, *P*(*E*|*C*), represents the test group whereas the second term, *P*(*E*|¬*C*), represents the control group. Δ*P* is the regression slope from *C* to *E*, letting

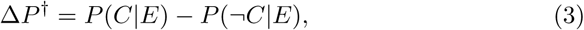

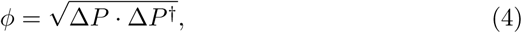

neglecting the standard deviations. Despite its rational status, Δ*P* does not fit the responses of humans in causal induction experiments well, and it is easy to design an experiment in which there is no correlation between the data and the prediction by Δ*P* predictions (e.g., see HO07.2 and W03.2 in the meta-analysis below, in Table 2). This index is at the core of most Bayesian models of causal induction (Cheng, 1997; Griffiths and Tenenbaum, 2005; Lu, Yuille, Liljeholm, Cheng and Holyoak, 2008).

**Table 2:**
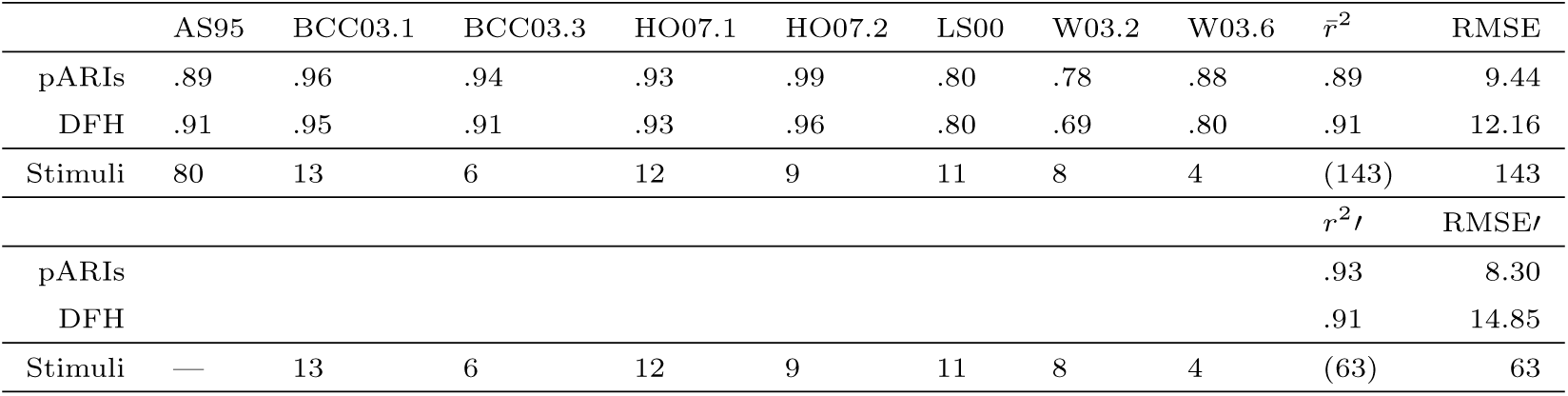
Replication of the meta-analysis in Hattori and Oaksford (2007), showing the determination coefficient *r*^2^ between the model predictions and the mean of the participants’ responses.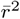 is the weighted average with Fisher’s *z*-transformation (the weights are the number of stimuli minus three), and RMSE is the root mean square error. 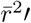 and RMSE′ are the results excluding the AS95 experiment. The last rows show the number of stimuli.

| | AS95 | BCC03.1 | BCC03.3 | HO07.1 | HO07.2 | LS00 | W03.2 | W03.6 | $\bar{r}^2$ | RMSE |
| --- | --- | --- | --- | --- | --- | --- | --- | --- | --- | --- |
| pARIs | .89 | .96 | .94 | .93 | .99 | .80 | .78 | .88 | .89 | 9.44 |
| DFH | .91 | .95 | .91 | .93 | .96 | .80 | .69 | .80 | .91 | 12.16 |
| Stimuli | 80 | 13 | 6 | 12 | 9 | 11 | 8 | 4 | (143) | 143 |
| | | | | | | | | | $r^2_{/}$ | RMSE$_{/}$ |
| pARIs |  |  |  |  |  |  |  |  | .93 | 8.30 |
| DFH |  |  |  |  |  |  |  |  | .91 | 14.85 |
| Stimuli | — | 13 | 6 | 12 | 9 | 11 | 8 | 4 | (63) | 63 |

DFH was introduced as a descriptive model, which is the geometric mean of the ‘predictability’ of effect from cause *P*(*E*|*C*) and its converse, the ‘diagnosability’ of cause from effect *P*(*C*|*E*), predicting that humans feel the strong causality only when both are high:

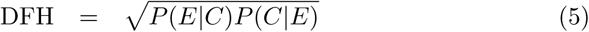

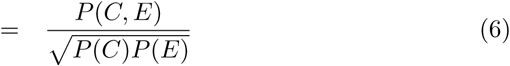

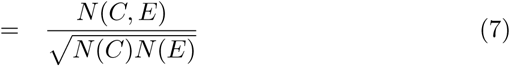

DFH showed the highest correlation with the data among models in the meta-analysis in Hattori and Oaksford (2007), compared with 32 other non-parametrised and 8 parametrised models. As grounds for the rationality of this model, Hattori and Oaksford showed that DFH is a limiting case of *ϕ* where *C* and *E* are both rare:

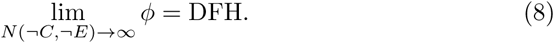

The limit operation is called an extreme rarity assumption; it neglects the assumed-to-be very abundant *N*(¬*C*, ¬*E*)-cell information.

Takahashi et al. (2010) have proposed the pARIs model, which is similar in some ways to DFH, but simpler:

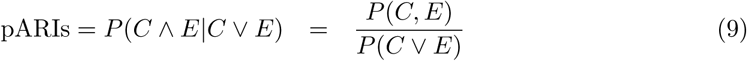

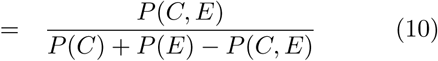

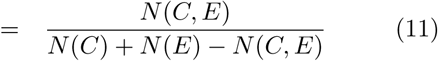

This model was named in reference to the proportion of confirmatory instances (pCI) model, proposed by White (2003), which is defined as:

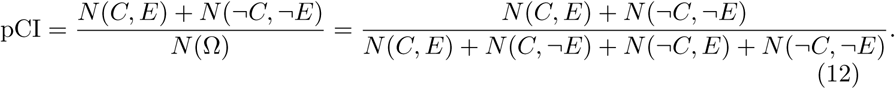

pARIs and pCI are conceptually different in that pARIs is a conditional probability and pCI is not (except in a trivial sense). Instead of the limit operation used in deriving DFH from *ϕ*, pARIs is based on the conditioning on the cases in which either *C* or *E* (or both) occurred. When events *C* and *E* are rare, among joint events between them, the three joint events, *C* ∧ *E, C* ∧ ¬*E* and ¬*C* ∧ *E* are rarer than the joint absence ¬*C* ∧ ¬*E*. The conditionalisation sets the focus on the rare joint events.

## 3. Computational analysis

According to the two-stage theory of causal induction—the distinctiveness of the stages has been empirically confirmed in (Hattori et al., 2017) —in the first, observational stage, the goal is to weed out unrelated event pairs and to sift a small number of correlated ones. Whether the correlated relationship is causal (and also the direction of the relationship) can be tested only in the second, interventional stage. Below, we show that DFH can be considered an index of dependence detection under an extreme rarity of causal events, and is rational in that sense, as far as it does relevance detection that is the principal goal of the first stage. The DFH model has been criticised as *non-normative* in that it is ‘not derived from a well-specified computational analysis of the goals of causal learning’ and is a groundless heuristic in (Lu et al., 2008). Here, we give a computational analysis of observational causal induction and rationalise DFH and pARIs.

In the analysis below, we heavily rely on rarity assumption. Rarity assumption in causal induction may be a rational prior as far as the causal events are formed similarly to natural categories that are based on family resemblance (Rosch, 1978; Navarro and Perfors, 2011; Gershman, 2019)^1^.

### 3.1. Statistical independence under extreme rarity assumptions

If *C* and *E* are statistically independent and *P*(*C*) and *P*(*E*) are non-zero, the following three terms are equal:

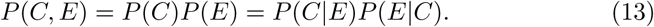

In terms of frequency, Eq. 13 is

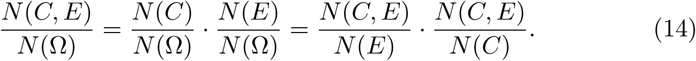

The derivation of DFH is based on an extreme rarity assumption represented by the limit operation where the occurrence probabilities of *C* and *E* are both very small. The rarity assumption is often used in probabilistic modelling of higher cognition (e.g., Oaksford and Chater (1994); McKenzie and Mikkelsen (2007)). In conditions of extreme rarity, i.e., *N*(¬*C*, ¬*E*) → ∞, although the first and second terms go to 0 because of the divergence of the denominator *N*(Ω), the third term (the product of conditional probabilities) stays invariant under the limit operation. Here, *P*(*C*|*E*)*P*(*E*|*C*) being close to 0 indicates the independence of *C* and *E*, whereas the term being close to 1 shows that *C* and *E* are dependent. The third term is the square of DFH, and

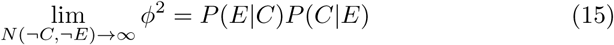

holds, which means that *P*(*C*|*E*)*P*(*E*|*C*) is the limiting case of the coefficient of determination on the 2×2 table, *ϕ*^2^. On the other hand, the second term of Δ*P*, *P*(*E*|¬*C*), vanishes and only *P*(*E*|*C*) remains: lim_*N*(¬*C*,¬*E*)→∞_ Δ*P* = *P*(*E*|*C*). *P*(*E*|*C*) fits the data quite poorly in the meta-analysis in (Hattori and Oaksford (2007), Model 12).

DFH is the square root of *P*(*C*|*E*)*P*(*E*|*C*), the measure for dependence under extreme rarity, and pARIs is its approximation within the range of single conditional probabilities.

### 3.2. pARIs and DFH under extreme rarity

Under extreme rarity, an index should take a certain value when *C* and *E* are independent, and its antipodal value when *C* and *E* are not independent, as discussed concerning Eq. 14. In this regard, when *C* and *E* are independent (see Eq. 13),

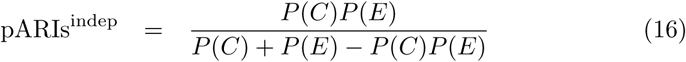

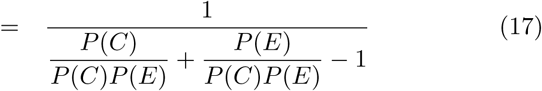

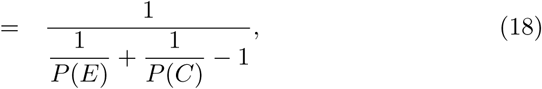

which implies that if *P*(*C*) and *P*(*E*) are both small (large), it goes to 0 (1). Also,

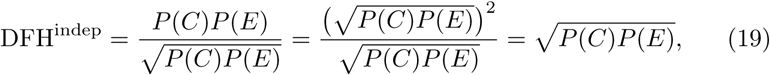

under independence, and shows a similar property to pARIs^indep^. We can also see that pARIs and DFH are 0 and 1 when *P*(*E*|*C*) = *P*(*C*|*E*) = 0 and *P*(*E*|*C*) = *P*(*C*|*E*) = 1, respectively. The difference in the behaviour of pARIs and that of DFH is how they connect the two extreme values, 0 and 1. It holds that pARIs ≤ DFH, because letting *U* = max{(*P*(*C*), *P*(*E*)}, *P*(*C*)*P*(*E*) ≤ *U* ^2^ and hence, 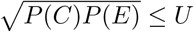. From Eqs. 6 and 10:

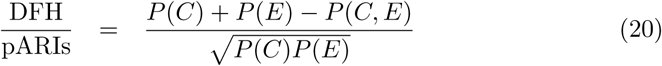

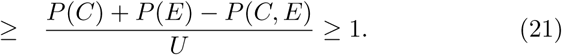

This means that pARIs is more prudent or conservative than DFH in that it takes a high value only when the independence is very high (Takahashi and Oyo, 2014), which could be useful in weeding out unrelated pairs in the first stage.

## 4. Meta-analysis

To compare the fit of the models introduced in the previous section, we reproduced the meta-analysis by Hattori and Oaksford (2007)^2^, with which they showed that DFH best fits the data among 41 existing indices. The experiments included in the meta-analysis were on causality, correlation or contingency, and selected according to the following five conditions. (1) The participants were humans, (2) there was only one effect and one candidate cause, (3) the answers were nearly continuous values, (4) the samples were given sequentially according to the predetermined frequencies, and (5) the causes were generative (a cause makes the effect occur), not preventive (a cause prevents the effect from occurring). In terms of the stimulus given to the participants, the last condition is defined by *N*(*C, E*)*N*(¬*C*, ¬*E*) ≥ *N*(*C*, ¬*E*)*N*(¬*C, E*), which is equivalent to *ϕ* ≥ 0 and Δ*P* ≥ 0.

Eight experiments are included in the meta-analysis: Experiment 1 of Anderson and Sheu (1995) (AS95); Experiments 1 and 3 of Buehner, Cheng and Clifford (2003) (BCC03.1 and BCC03.3)^3^; Experiments 1 and 2 of Hattori and Oaksford (2007) (HO07.1 and HO07.2); Experiments 1–3 of Lober and Shanks (2000) (LS00); and Experiments 2 and 6 of White (2003) (W03.2 and W03.6). To measure each index’s fit to the data, we calculated the coefficient of correlation (*r*) between each index and participants’ mean ratings of causal strength.

Among the eight experiments included, AS95 differs from others in some ways. One is that the number of stimuli is very large (80 out of 143, the total of all eight experiments: more than the half of the total), which gives a much higher weight to AS95 in calculating the overall result than all other experiments. In this experiment, 40 participants responded to all the 80 stimuli, taking one hour, which is much longer than for other experiments, and the stimuli varied differentially. Although the participants should independently respond to each stimulus, totally forgetting the past stimuli presented before, it is possible that they kept learning co-occurrence information; hence, their responses may have become increasingly confounded with past stimuli as the experiment proceeds. Although this effect may have happened in other experiments, it may be the most prominent in AS95.

Even without the possible problems mentioned above, it is not ideal that a meta-analysis result depends so much on a single experiment that necessarily has a specific set of conditions. Therefore, we calculated the overall data fit with and without AS95, as shown in Table 2.

### 4.1. Result

In replicating the meta-analysis, we completely reproduced the results in Hattori and Oaksford (2007), except for the small mistake in the original data as mentioned in Footnote 1. As we confirmed that DFH shows the best fit among the 33 non-parametrised models, we focused on comparing DFH and pARIs.

Table 2 shows the result of the meta-analysis of DFH and pARIs. Despite its simple form, pARIs has the same high correlation to the data in comparison with DFH, with the lowest fit of *r*^2^ = .78 to W03.2, and *r*^2^ = .69 for DFH. On the data from the AS95 experiment, which has the largest number of responses (80), DFH fit better than pARIs. We show the overall fit and error for the seven experiments (excluding AS95) in Table 2 as 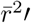 and RMSE′, as well. Without AS95, pARIs fits the data better than DFH, and vice versa with AS95. The root mean squared error for pARIs is overall smaller than DFH, with or without AS95.

### 4.2. Discussion

We conclude that pARIs has the same level of descriptive performance as DFH in many cases, despite its simpler definition. pARIs fits the data as well as DFH, which means that the data fit of pARIs is also better than the other 32 non-parametrised and 8 parametrised models compared in Hattori and Oaksford (2007).

From the result of the meta-analysis, we can state that the causal intuition of human beings roughly follows the pARIs rule, as well as that of DFH. Then, what are the adaptive merits of having a pARIs-like causal intuition? One way to answer the question is computer simulations as carried out in Hattori and Oaksford (2007).

## 5. Critical examination of the simulations

Hattori and Oaksford ran simulations that showed the adaptive merit of DFH-like causal intuition which is that it approximates the sample and population correlation coefficients *ϕ* and *ϕ*_0_ well from contingency tables under some realistic conditions, such as limitations to working memory and the rarity and equiprobability of cause and effect (Hattori and Oaksford, 2007). It is considered the main part of their rational analysis (Anderson, 1990) which is to test the optimality of a part of cognition in terms of the cognitive limitations of the agent and the structure of the environment in which the cognitive agent acts and adapts.

In their paper, Simulation 1 tests effectiveness and parsimony (Fig. 3a, 3b, and 4 in Hattori and Oaksford (2007)) of some models and Simulation 2 treats their efficiency (Fig. 5 in Hattori and Oaksford (2007)). For convenience, we rearrange the simulations into three groups rather than two. Simulation A (Fig. 3a in Hattori and Oaksford (2007)) tests descriptive performance of DFH, that is, how similarly it behaves to *ϕ* (the correlation coefficient of the sample). Simulation B (Fig. 3b and 4 in Hattori and Oaksford (2007)) tests inferential performance, or how well DFH and *ϕ* can be used to infer the value of *ϕ*_0_, the correlation coefficient of the population. Simulation C (Fig. 5 in Hattori and Oaksford (2007)) tests the speed and accuracy of the inference process as DFH sequentially approximates the value of *ϕ*_0_ and shows the time development of the index values.

Throughout the simulations, there are two kinds of sampling. One is the usual sampling (‘*N* -sampling’) where the number of instances is *N* = *N*(*ω*) = *N*(*C, E*) + *N*(*C*, ¬*E*) + *N*(¬*C, E*) + *N*(¬*C*, ¬*E*). The other is called ‘*N*_*W*_ - sampling’ where *N*_*W*_ = *N*(*C* ∨ *E*) = *N*(*C, E*) + *N*(*C*, ¬*E*) + *N*(¬*C, E*). In the latter sampling, inspired by consideration of limitation in working memory capacity ^4^, only the assumed-to-be rare instances are counted, and the number of assumed-to-be abundant instances, *N*(¬*C*, ¬*E*), is neglected. For example, it can be that *N*_*W*_ = 7 while *N* = 10 when *N*(¬*C*, ¬*E*) = 3. In Simulation C, the time development of the simluation ticks with the increments of *N* or *N*_*W*_. See Takahashi and Oyo (2014) for the preliminary results, including those pARIs. Here, we just point out two implicit problems in the original simulations.

### 5.1. Problem 1: Samples being always generative

The first problem is the implicit filtering of the contingency tables generated from the probability distribution. The probability distributions in the simulations were fixed to be *generative* (*C* and *E* are positively correlated, i.e. *P*(*C, E*) ≥ *P*(*C*)*P*(*E*)) rather than *preventive* (*C* and *E* are negatively correlated, i.e. *P*(*C, E*) *< P* (*C*)*P*(*E*)), because of the scope of their theory and model. The problem is that the contingency tables used for calculating and evaluating the indices are limited to generative ones, i.e., where *N*(*C, E*)*N*(¬*C*, ¬*E*) *> N* (*C*, ¬*E*)*N*(¬*C, E*) holds. We have confirmed this problem with an exhaustive test of possible simulation settings. The probability distribution being generative does not mean that its sample is always generative. This implicit condition of limiting the samples to be generative was imposed on Simulation A and B, but not C.

We think that preventive tables sampled from generative populations should be included, because it is unnatural that agents can discard contingently preventive cases and keep sampling until the sample is generative with the same, fixed generative population. Preventive samples should be judged at a lower value of the indices, rather than arbitrarily discarded. In preliminary simulations, we found that the correlation between DFH and *ϕ* is generally lower when preventive tables are included, when *P*(*C*) and *P*(*E*) are not small. Seemingly, it does not affect the interpretations of the original result that focus on the cases in which *P*(*C*) and *P*(*E*) are both small, but further investigation is needed.

### 5.2. Problem 2: Definability of indices on the samples

The other problem is in the treatment of contingency tables on which some indices are not defined. For example, DFH takes the undefined value 0*/*0 because of the zero denominator, for example when (*N*(*C, E*), *N*(*C*, ¬*E*), *N*(¬*C, E*), *N*(¬*C*, ¬*E*)) = (0, 0, 1, 10), whereas pARIs takes the value 0. The cases in which the indices are defined (i.e. have a definite value) have an inclusion order, and when *ϕ* is defined, DFH is always defined, as we see below. The possible undefinedness affects the significance of all three simulations, especially the early stage of Simulation C.

We analyse the ‘definability’ of the models, especially because we deal with very small samples that model the quite limited capacity of our working memory, and we often do not have sufficient time to gather data, nor to wait for more occurrences. The models have different definability, as summarised in Table 3, where the combinations of the values of three cells—excluding *N*(¬*C*, ¬*E*), zero or positive—are considered. In the eight cases, min *N* ≤ 4 and min *N*_*W*_ ≤ 3. As for the four indices *ϕ*, Δ*P*, DFH and pARIs, the more complicated the index is, the fewer cases it is defined for. *ϕ*, Δ*P*, DFH and pARIs have 5, 6, 5 and 7 definable cases, respectively. pARIs is defined simply whenever *N*_*W*_ ≥ 1. DFH is defined when min *N*_*W*_ ≥ 2 or *N*(*C, E*) ≥ 1. A sufficient condition for *ϕ* and Δ*P* to be defined is min*N* ≥ 3.

**Table 3:**
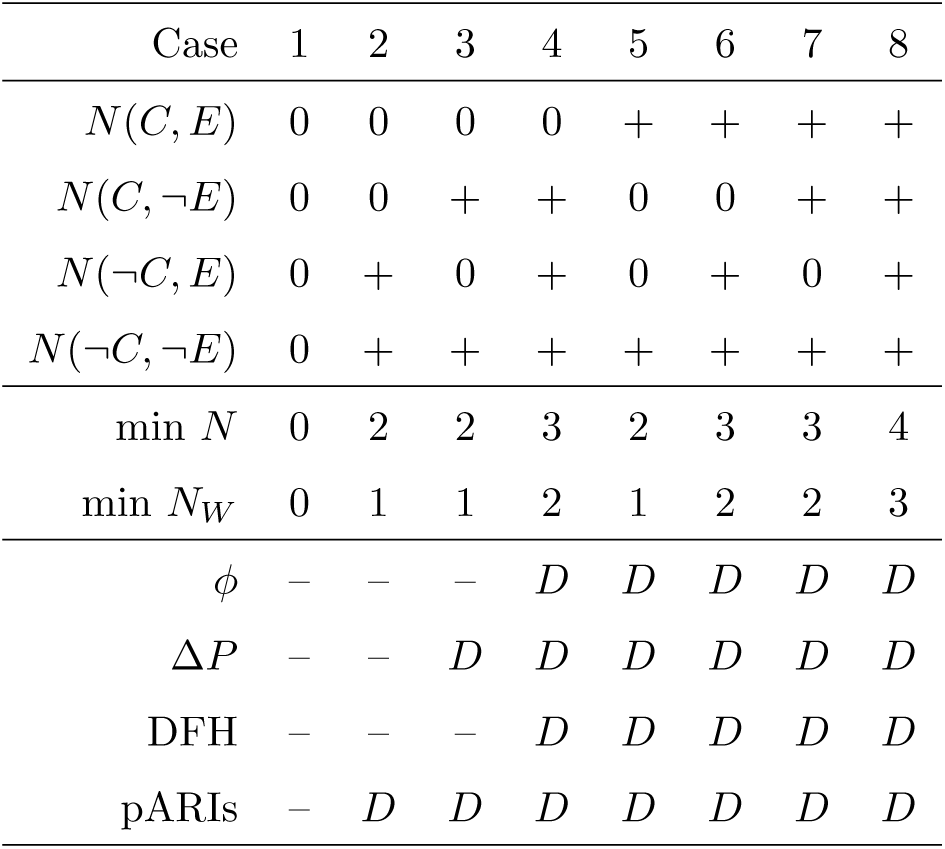
The definability (*D*: defined, –: not defined) of the four indices *ϕ*, Δ*P*, DFH and pARIs, on the 2^3^ = 8 cases where three frequencies *N*(*C, E*), *N*(*C*, ¬*E*) and *N*(¬*C, E*) are zero or positive (+). We fixed *N*(¬*C*, ¬*E*) to be positive in *N*_*W*_ ≥ 1 cases for simplicity.

| Case | 1 | 2 | 3 | 4 | 5 | 6 | 7 | 8 |
| --- | --- | --- | --- | --- | --- | --- | --- | --- |
| $N(C, E)$ | 0 | 0 | 0 | 0 | + | + | + | + |
| $N(C, \neg E)$ | 0 | 0 | + | + | 0 | 0 | + | + |
| $N(\neg C, E)$ | 0 | + | 0 | + | 0 | + | 0 | + |
| $N(\neg C, \neg E)$ | 0 | + | + | + | + | + | + | + |
| $\min N$ | 0 | 2 | 2 | 3 | 2 | 3 | 3 | 4 |
| $\min N_W$ | 0 | 1 | 1 | 2 | 1 | 2 | 2 | 3 |
| $\phi$ | - | - | - | $D$ | $D$ | $D$ | $D$ | $D$ |
| $\Delta P$ | - | - | $D$ | $D$ | $D$ | $D$ | $D$ | $D$ |
| DFH | - | - | - | $D$ | $D$ | $D$ | $D$ | $D$ |
| pARIs | - | $D$ | $D$ | $D$ | $D$ | $D$ | $D$ | $D$ |

The original simulations in Hattori and Oaksford (2007) did not consider definability. Definability can affect the validity of the results if indices with different definability are compared, and this implicitly happened in their study. A valid comparison would be to say one model is better than another only when the former is defined on strictly more cases than the latter and still performs better.

In Simulation A (Fig. 3a of Hattori and Oaksford (2007)), only contingency tables in which *ϕ* is defined are included in the result. When the sample size is small (around *N* = 7), a large part of the contingency tables with small *P*(*C*) and *P*(*E*) do not define *ϕ*: only Cases 4–8 in Table 3 do. It discounts the meaningfulness of Simulation A in terms of small samples, and this is a heavy condition to impose on the tables when *N* or *N*_*W*_ is so small such as *N* ∼ 𝒩 (7, 1^2^), especially when we assume that *P*(*C*) and *P*(*E*) are both small. We calculated the proportion of tables generated in Simulation A on which DFH is defined, which were around 35% and 65% of the sampled tables when *P*(*C*) = *P*(*E*) = .1 and *P*(*C*) = *P*(*E*) = .2, respectively. pARIs is defined on around 62% and 86% of the tables, respectively, which means that pARIs faces this problem strictly less than DFH.

This condition of *ϕ* being definable is also imposed on Simulation B, although it is not necessary in this simulation setting because *ϕ*_0_ is the measure to be inferred. pARIs gives qualitatively similar but significantly better inferences of the population correlation coefficient *ϕ*_0_ in Simulation B than DFH, even though pARIs is computed on a wider variety of contingency tables.

In Fig. 5 in Hattori and Oaksford (2007) from Simulation C, in which the time developments of the mean index value of the indices from sample sizes 1 to 30 are shown, the mean value of DFH always decreases from 1.0 at sample size 1, because DFH is definable only for (*N*(*C, E*), *N*(*C*, ¬*E*), *N*(¬*C, E*)) = (1, 0, 0) when *N* = 1 or *N*_*W*_ = 1. Such samples are less than 20% at *N* = 1, and 18%–90% at *N*_*W*_ = 1, depending on the population correlation coefficient *ϕ*_0_ ∈ {.1, .2, …, .9}. The percentages monotonically increase in *N* or *N*_*W*_ increments. The percentages of the samples on which pARIs can be defined are 20%–40% at *N* = 1 and 100% at *N*_*W*_ = 1, and a strict order relation holds with DFH, as we can see in Table 3.

### 5.3. Summary

Although DFH is the limiting case of the correlation coefficient, pARIs better infers the population correlation, and in much wider cases. This property makes a larger difference when considering multiple causes and effects, making the simulation settings more realistic.

## 6. General discussion

### 6.1. DFH and pARIs

We begin our discussion by comparing pARIs and DFH, the two similar models. Both are related to causal biconditionals, (i.e. ‘if cause then effect, and if effect then cause’). DFH represents both conditionals as the geometric mean of *P*(*E*|*C*) and *P*(*C*|*E*), and pARIs directly uses the probability of the biconditional:

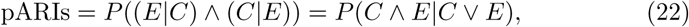

according to representative theories of compound conditionals (McGee, 1989; Kaufmann, 2009; Gilio and Sanfilippo, 2014; Sanfilippo, Pfeifer, Over and Gilio, 2018), or in a conditional event algebra initiated in de Finetti (1964) and then developed in, e.g., Goodman, Nguyen and Walker (1991). In the formal systems, *E*|*C* and *C*|*E* are themselves considered to be events, and hence can be logically connected.

Another difference between pARIs and DFH is the process by which they ignore the absence-absence, ¬*C* ∧ ¬*E* cases. pARIs conditions on *C* ∨ *E* cases, whereas DFH starts from *ϕ* and assumes that *N*(¬*C*, ¬*E*) diverges to positive infinity. Let us consider the nature of ¬*C* ∧ ¬*E* cases to see which is more appropriate for dealing with them.

The prerequisite for frequency information is counting occurrences. However, how do we count the absence, or non-occurrence, of events? Often, counting non-occurrences is indefinite or ill-defined. When we consider the causal relationship between the effect of an engine starting (*E*) and a candidate cause of turning the key (*C*), there are four cases of co-occurrence. We have no difficulty in counting the other three cases, but the ¬*C* ∧ ¬*E* cases are problematic. How many times have we observed ‘no key turning (¬*C*) and no engine starting (¬*E*)’? We could say we have observed this case infinite times (*N*(¬*C*, ¬*E*) → ∞). We can also say that we have observed it as many as the intervals between key turning and/or engine starting (*N*(¬*C*, ¬*E*) := *N*(*C* ∨ *E*) − 1), or we might want to say we have never observed such an event.

The count of *N*(¬*C*, ¬*E*) depends on the spatio-temporal framing of events. A more definite framing than the examples above is the division of a day into 144 ten-minute intervals and the identification of occurrences of *C* and *E*. Or we can count the occurrence of times when a driver enters a car, turns the key, and then exits the car after we observe the effect. In this way, ¬*C* ∧ ¬*E* cases can be made definite, although there are many ways to frame it. In our everyday causal judgment, it would be quite rare for framing to be rigorous and unambiguous. Defining the unit for ¬*C* ∧ ¬*E* cases is a necessary condition for determining the *probabilistic model* in the sense of assignment of probabilities to atomic events. In other words, the probabilistic models that we use in everyday life are often not well-defined. In contrast, modern scientific experiments with a control group have a well-defined framing. The division of test and control groups itself is the effect of intervention, and causal induction is no longer purely observational.

Considering the indefiniteness of framing in everyday causal induction, in which we can count ¬*C* ∧ ¬*E* cases arbitrarily, neglecting *N*(¬*C*, ¬*E*) by conditioning may be more natural and robust than always expecting countless *N*(¬*C*, ¬*E*). pARIs can be considered conditioning on *C* ∨ *E* of the conjunctive case *C* ∧ *E* or of the probability of logical equivalence *P*(*C* ∧ *E*) + *P*(¬*C* ∧ ¬*E*) (equivalent to the pCI rule). Considering the often indefinite *N*(¬*C*, ¬*E*), ignoring them via conditioning and judgment about the statistical relationship between *C* and *E* from the three other, more definite cases would give a stable estimate. pARIs performs precisly this function.

### 6.2. Causal induction models in terms of conditionals

We compare the three models pARIs, DFH and Δ*P*, in terms of their use of conditionals, based on the fact that the subjective probability of a causal conditional ‘if *C*, then *E*’ is well described by the conditional probability *P*(*E*|*C*) (e.g., see Over, Hadjichristidis, Evans, Handley and Sloman (2007), but also see Skovgaard-Olsen, Singmann and Klauer (2016) for criticism and generalization).

Δ*P* = *P*(*E*|*C*)− *P*(*E*|¬*C*) = *P*(*E*|*C*)+*P*(¬*E*|¬*C*)−1 is the adjusted sum of the probabilities of the conditional ‘if *C*, then *E*’ and its inverse, ‘if not *C*, then not *E*’, which correspond to sufficient and necessary conditions, respectively (Pearl, 2000). However, the latter probability vanishes under extreme rarity. In pARIs and DFH, the converse (‘if effect then cause’, *P*(*C*|*E*)) is used as a kind of necessary condition, instead of the inverse, which stays valid under the rarity assumption. Although the contrapositive conditionals, such as the inverse (‘if not cause then not effect’) and the converse (‘if effect then cause’), are equivalent in classical logic, they are not equivalent in probabilistic logic. Still, *P*(*C*|*E*) = *N*(*C, E*)*/N* (*E*) and *P*(¬*E*|¬*C*) = *N*(¬*C*, ¬*E*)*/N* (¬*C*) have the common element *N*(¬*C, E*) in their denominators, and therefore when *N*(*C, E*) and *N*(¬*C*, ¬*E*) are close, or if the value of *N*(*C, E*) is used in place of *N*(¬*C*, ¬*E*), especially when the latter is indefinite or uncertain, *P*(*C*|*E*) could well work on behalf of *P*(¬*E*|¬*C*).

Several authors have theoretically shown that biconditionals of the form ‘if *p*, then *q*, and if *q*, then *p*’ has the form similar to pARIs as in Eq. 22 (McGee, 1989; Kaufmann, 2009; Gilio and Sanfilippo, 2014; Sanfilippo et al., 2018). Baratgin, Politzer, Over and Takahashi (2018) have confirmed that the abstract (non-causal) biconditionals has a well corresponding truth table to pARIs. In the subjective probability theory developed by De Finetti, conditional probabilities are atomic and that they correspond to conditional events (de Finetti, 1964).

Causal biconditionals and pARIs-like causal intuition likely have a more direct relationship, as suggested in Gauffroy and Barrouillet (2009). The causal relationship between cause and effect may be recognised when both directions of conditional—’if cause, then effect’ (predictive) and ‘if effect, then cause’ (diagnostic)—are acceptable (i.e., *P*(effect|cause) and *P*(cause|effect) are both high). Considering only the predictive conditional is sufficient for predicting the occurrence of effect from that of cause, but it does not exclude the possibility that *P* (effect|cause) is high just because *P*(effect) is high and cause is not strongly preventive of effect. Evaluating *P*(cause|effect) at the same time may be an effective strategy for considering the necessity of the cause for the effect.

In a Bayesian framework, Lu et al. point out that people favour sparse and strong causes (Lu et al., 2008). If we interpret a cause *C* as being strong when *P*(*E*|*C*) is high, and sparse when *P*(*C*|*E*) is high, it can be modelled as a high biconditional probability. One situation in which sparseness can be modelled as *P*(*C*|*E*) being high is when multiple causes *C*_1_, *C*_2_, …are mutually exclusive. *P*(*C*_*i*_|*E*) being high means that all *P*(*C*_*j* ≠*i*_|*E*) are low. In this sense, the form of pARIs can be considered a simple expression of people’s priors on causal relationships.

### 6.3. Computational cost of causal induction models

Forming a conditional probability would elicit some cognitive cost through the Ramsey test (Ramsey, 1990) or alike. Two conditional probabilities are required to compute Δ*P* and DFH, whereas pARIs needs a single conditional probability. Still, pARIs accounts for two reciprocal conditionals, consistent with the general tendency that single (rather than plural) hypotheses are preferred (e.g. the singularity principle by Evans (2007)). More complicated or computationally demanding Bayesian models (e.g., causal support in Griffiths and Tenenbaum (2005) and the SS model in Lu et al. (2008)) are not excluded as cognitive models. However, the kind of Bayesian computations that can realistically or effectively be approximated in our brains must be investigated further.

In all cases, pARIs is advantageous as a cognitive model because it is a simple, single conditional probability that integrates properties like those of Δ*P* and DFH. This makes a bigger difference when we consider many-to-one or many-to-many cause-effect correspondences, which are the computational goal of causal search to be done in the observational stage of causal induction.

### 6.4. Observational causal induction as similarity judgment

As we have seen in the previous section, there are several indices of similarity and other concepts equivalent to pARIs. Moreover, DFH is also equivalent to a similarity indices called cosine similarity (or the Ochiai similarity index in Ochiai (1957)). It is mathematically interesting that the correlation coefficient *ϕ* ∈ [−1, 1] becomes the similarity index DFH ∈ [0, 1] under the limit operation, as in Eq. 8 (equation (2) in Hattori and Oaksford (2007)). pARIs is also equivalent to the Jaccard similarity index (Jaccard, 1901), which is commonly used in statistical analysis including cluster analysis in biology and sociology (e.g., Kaufman and Rousseeuw (1990)), and gives rise to a proper distance metric between sample sets (Gilbert, 1972). These facts suggest that humans judge statistical similarity between the occurrences of *C* and *E* when they observe the statistical information to verify a correlation between the two events. It is consistent with the hypothesis that Chater and Oaksford (2006) proposed: the human ability to learn causal relationships remarkably quickly—sometimes even in a single instance—depends on the feature similarity of the candidate cause and the effect.

Succeeding the study by Hattori and Oaksford (2007), this study suggested that people respond with statistical similarity to ascertain the strength of causal relationships in a realistic setting with a small sample and no other cues (spatial or temporal information, prior knowledge, or causal models). According to premilinary simulation results in Takahashi and Oyo (2014), similarity indices— especially pARIs—contributes to fast and accurate inference of correlation under the rarity assumption, at least when compared with correlation (*ϕ*) and regression (Δ*P*) indices^5^.

### 6.5. Future work

Establishing observational causal induction as a stastical independence test, and pARIs as the appropriate model will require further theoretical and empirical arguments. To expand the theoretical aspect, we must situate the observational stage into a more comprehensive framework of causal inference. One possibility is a causal search in which the topology (graph structure) of a causal Bayes network, rather than the edge weights (strength), is established (Griffiths and Tenenbaum, 2005; Pearl, 2000). In the PC algorithm for causal search (Spirtes and Glymour, 1991), which is a representative constraint-based algorithm for constructing sparse causal graphs, the complete undirected graph of all the variables is formed, and then undirected edges are removed if they are (conditionally) independent. The directionality of the edges between nodes is determined only after the removal of undirected edges. Our hypothesis is that people use pARIs- or DFH-like intuition to quickly test the independence quickly under assumptions like rarity of events, explaining why DFH and pARIs are symmetric or undirected (i.e. invariant under exchange of cause and effect), and may resolve some of the rationality issues posed in Lu et al. (2008).

Empirically, we can investigate whether people are trying to test independence, and when they do so, which factors contribute to switching behaviours and goals, as in Hattori et al. (2017). To compare DFH and pARIs, we can set the stimuli so that the predictions by the two models diverge. DFH depends on the value of *N*(*C*, ¬*E*) × *N*(¬*C, E*) whereas pARIs does not, so we can keep *N*(*C*, ¬*E*) + *N*(¬*C, E*) invariant and vary *N*(*C*, ¬*E*) × *N*(¬*C, E*).

## 7. Conclusion

Adopting the framework of the two-stage theory of human causal judgment Hattori and Oaksford (2007), we have proposed a computational analysis of the first, observational stage of causal induction. According to this analysis, when an extreme rarity assumption is valid, it is rational to use the product of the two reciprocal conditional probabilities of two events to detect non-independence between them. The DFH model is the square root of this measure, and the pARIs rule effectively approximates it. We replicated the meta-analysis in Hattori and Oaksford (2007) and showed that DFH and pARIs both fit the data well, even though the latter is simpler. We then pointed out two problems in the simulations as a rational analysis of causal induction in Hattori and Oaksford (2007). We showed that a crucial problem can be solved if we adopt pARIs as the model derived from the theory. The difference between DFH and pARIs is, as far as the two models represent causal biconditionals between cause and effect, that the former is merely the geometric mean of the two conditional probabilities, whereas the latter has rational grounding as the probability of a biconditional event. Adopting the rarity assumption, which has been proven important in covariation assessment (McKenzie and Mikkelsen, 2007) and other reasoning tasks (e.g. Oaksford and Chater (1994)), we conducted a new computational analysis and proposed a new model consistent with the theories of compound conditionals that further supports the new paradigm psychology of reasoning advocated by David Over (Over, 2009).

## 8. Acknowledgments

We thank David Over for his generous support. We also thank Jean Baratgin, Mike Oaksford, and Nicole Cruz for helpful discussions and suggestions, and Masasi Hattori for providing us with the raw data of the meta-analysis. Part of this work was supported by a Grant-in-Aid for Scientific Research 17H04696 from the Japan Society for the Promotion of Science.

## Footnotes

1 Interestingly, normalizing and linearly transforming the goodness of a category formed based on family resemblance, Eq. (7) in Navarro and Perfors (2011), gives an equivalent index as pARIs, which reinforces our argument in 6.4.

2 There was a small mistake in the meta-analysis in Hattori and Oaks-ford (2007). The mean of the responses by the participants for the stimulus (*N*(*C, E*), *N*(*C*, ¬*E*), *N*(¬*C, E*), *N*(¬*C*, ¬*E*)) = (28, 0, 21, 7) in the experiment abbreviated as LS00 (in Experiment 2 in Lober and Shanks (2000)). It was listed as 34 in the analysis, but is actually 65. This correction does not affect the result very much and therefore their argument is not affected.

3 In the BCC03.1 experiment, there are 15 stimuli, of which only 13 are handled in Hattori and Oaksford (2007). We excluded two stimuli where *P*(*E*|*C*) = *P*(*E*|¬*C*) = 0 and *P*(*E*|*C*) = *P*(*E*|¬*C*) = 1 and roughly replicated the original result. For the former, DFH and the Power PC model (Cheng, 1997) cannot be computed because their denominators are zero (i.e. Case 3 in Table 3). We guess that the latter was excluded because the stimulus was not strictly generative.

4 In Hattori and Oaksford (2007), the limitation in working memory capacity is argued in terms of the magic number 7 ± 2, which has recently been considered to be around 4 (Cowan, 2001). However, especially considering the small samples treated in their and our studies, where the numbers on all four cells tend to be small (all usually one digit in the decimal system), it could be more appropriate to consider the number of joint event types, instead of the total frequencies, *N* or *N*_*W*_, to be within the limitation. One prediction that can be derived from this is that people would quite accurately remember the total frequency of four types such as on the 2 × 2 contingency tables, while people would tend to ignore some of the joint event types on the 3 × 3 or larger contingency tables. The 3 × 3 tables are called for in cases in which some of the occurrences of *C* or *E* may be uncertain, which often happens in scientific data collected by sensors (Yokokawa and Takahashi, 2014; Tanaka, Namiki, Oyo and Takahashi, 2015), and are similar to 3 × 3 de Finetti truth tables (Baratgin, Over and Politzer, 2013). Still, sample sizes as small as 7 ± 2 is a realistic condition that people can deal with only a smaller number of frequencies because of a temporal rather than memory limit, where we usually make an action as soon as possible, in addition to the memory limitation.

5 A reviewer pointed out that when *C* ∩*E* = ∅ (i.e. *N*(*C, E*) = 0), pARIs = 0, while Δ*P >* 0, and it is problematic for pARIs that it cannot express such negative probabilistic dependency. We think the comment might in principle extend to criticism of the framework of the present computational analysis, but we try to answer it approximately within the framework, from two aspects. First, Hattori and Oaksford (2007) discussed DFH (and hence pARIs) can deal with the negative probabilistic dependencies, or preventive causes, by transforming ‘*C* prevents *E*’ into ‘¬*C* causes *E*’, swapping the two rows on the contingency table. While Lu et al. (2008) criticized the idea, there is no empirical test done so far. Second, under the assumptions of our analysis, the goal is to compute the non-independence of *C* and *E* that is unipolar (non-negative). Also, as discussed in 6.2, under the assumptions, the second term of Δ*P* vanishes and merely Δ*P* = *P*(*E*|*C*). Even outside the assumptions, as discussed in 6.1, the value of *N*(¬*C*, ¬*E*) may be quite unreliable, and then Δ*P* may not work reliably. A related empirical work is Hattori et al. (2017) where they show that depending on some factors that directly affects the reliability of *N*(¬*C*, ¬*E*), people’s causal judgments vary, in some cases close to DFH, and in other cases to Δ*P*, which may be considered rational in the above sense.

## References

Anderson, J.R., 1990. The Adaptive Character of Thought. Hillsdele, NJ: Erlbaum.

Anderson, J.R., Sheu, C.F., 1995. Causal inferences as perceptual judgements. Memory & cognition 23, 510–24.

Baratgin, J., Over, D.E., Politzer, G., 2013. Uncertainty and the de Finetti tables. Thinking & Reasoning 19, 308–328.

Baratgin, J., Politzer, G., Over, D.E., Takahashi, T., 2018. The psychology of uncertainty and three-valued truth tables. Frontiers in Psychology 9, 1479. URL: https://www.frontiersin.org/article/10.3389/fpsyg.2018.01479, doi:10.3389/fpsyg.2018.01479.

Buehner, M.J., Cheng, P.W., Clifford, D., 2003. From covariation to causation: a test of the assumption of causal power. Journal of experimental psychology. Learning, memory, and cognition 29, 1119–1140.

Chater, N., Oaksford, M., 2006. Mental Mechanisms: Speculations on Human Causal Learning and Reasoning, in: Fiedler, K., Juslin, P. (Eds.), Information Sampling and Adaptive Cognition. Cambridge University Press, pp. 210–236.

Cheng, P.W., 1997. From covariation to causation: A causal power theory. Psychological Review 104, 367–405.

Cowan, N., 2001. The magical number 4 in short term memory. A reconsideration of storage capacity. Behavioral and Brain Sciences 24, 87–186. 0140-525X.

Evans, J.S.B.T., 2007. Hypothetical Thinking: Dual Processes in Reasoning and Judgement. Psychology Press.

de Finetti, B., 1964. Foresight: Its logical laws, its subjective sources (original publication, 1937), in: Kyburg, H., Smokier, H.E. (Eds.), Studies in subjective probability. Wiley, New York, pp. 55–118.

Gauffroy, C., Barrouillet, P., 2009. Heuristic and analytic processes in mental models for conditionals: An integrative developmental theory. Developmental Review 29, 249–282.

Gershman, S.J., 2019. How to never be wrong. Psychonomic Bulletin and Review 26, 13–28. doi:10.3758/s13423-018-1488-8.

Gilbert, G., 1972. Distance between Sets. Nature 239, 174–174.

Gilio, A., Sanfilippo, G., 2014. Conditional Random Quantities and Compounds of Conditionals. Studia Logica 102, 709–729. 1304.4990v1.

Goodman, I.R., Nguyen, H.T., Walker, E.A., 1991. Conditional Inference and Logic for Intelligent Systems: A Theory of Measure-Free Conditioning. Amsterdam: North-Holland.

Griffiths, T.L., Tenenbaum, J.B., 2005. Structure and strength in causal induction. Cognitive Psychology 51, 334–84.

Griffiths, T.L., Tenenbaum, J.B., 2009. Theory-based causal induction. Psychological Review 116, 661–716.

Gunji, Y.P., Takahashi, T., Aono, M., 2004. Dynamical infomorphism: Form of endo-perspective. Chaos, Solitons and Fractals 22, 1077–1101.

Hattori, I., Hattori, M., Over, D.E., Takahashi, T., Baratgin, J., 2017. Dual frames for causal induction: the normative and the heuristic. Thinking & Reasoning 23, 292–317.

Hattori, M., Oaksford, M., 2007. Adaptive Non-Interventional Heuristics for Covariation Detection in Causal Induction: Model Comparison and Rational Analysis. Cognitive Science 31, 765–814.

Hattori, M., Over, D.E., Hattori, I., Takahashi, T., Baratgin, J., 2016. Dual frames in causal reasoning and other types of thinking. The Thinking Mind: A Festschrift for Ken Manktelow, 98–114.

Jaccard, P., 1901. Étude comparative de la distribution florale dans une portion des Alpes et des Jura. Bulletin de la Société Vaudoise des Sciences Naturelles 37, 547–579.

Jenkins, H.M., Ward, W.C., 1965. Judgment of Contingency between Responses and Outcomes. Psychological Monographs: General and Applied 79, 1–17.

Kaufman, L., Rousseeuw, P.J., 1990. Finding Groups in Data: An Introduction to Cluster Analysis.

Kaufmann, S., 2009. Conditionals Right and Left: Probabilities for the Whole Family. Journal of Philosophical Logic 38, 1–53.

Lake, B.M., Ullman, T.D., Tenenbaum, J.B., Gershman, S.J., 2017. Building Machines That Learn and Think Like People. Behavioral and Brain Sciences 40, 1–72.

Lober, K., Shanks, D.R., 2000. Is causal induction based on causal power? Critique of Cheng (1997). Psychological review 107, 195–212.

Lu, H., Yuille, A.L., Liljeholm, M., Cheng, P.W., Holyoak, K.J., 2008. Bayesian Generic Priors for Causal Learning. Psychological Review 115, 955–984.

McGee, V., 1989. Conditional Probabilities and Compounds of Conditionals. The Philosophical Review 98, 485–541.

McKenzie, C.R.M., Mikkelsen, L.A., 2007. A Bayesian view of covariation assessment. Cognitive Psychology 54, 33–61.

Navarro, D.J., Perfors, A.F., 2011. Hypothesis Generation, Sparse Categories, and the Positive Test Strategy. Psychological Review 118, 120–134. doi:10.1037/a0021110.

Oaksford, M., Chater, N., 1994. A Rational Analysis of the Selection Task as Optimal Data Selection. Psychological Review 101, 608–631.

Ochiai, A., 1957. Zoogeographic studies on the soleoid fishes found in Japan and its neighbouring regions. Bull. Jpn. Soc. Sci. Fish 22, 526–530.

Over, D.E., 2009. New paradigm psychology of reasoning. Thinking & Reasoning 15, 431–438.

Over, D.E., Hadjichristidis, C., Evans, J.S.B.T., Handley, S.J., Sloman, S.a., 2007. The probability of causal conditionals. Cognitive psychology 54, 62–97.

Pearl, J., 2000. Causality: Models, Reasoning, and Inference. Cambridge University Press, New York, NY, USA.

Ramsey, F.P., 1990. General Propositions and Causality, in: Mellor, H.A. (Ed.), Philosophical Papers. Cambridge University Press.

Rosch, E., 1978. Principles of Categorization, in: Rosch, E., Lloyd, B.B. (Eds.), Cognition and categorization. Hillsdale, NJ: Lawrence Erlbaum, pp. 27–48.

Sanfilippo, G., Pfeifer, N., Over, D.E., Gilio, A., 2018. Probabilistic inferences from conjoined to iterated conditionals. International Journal of Approximate Reasoning 93, 103–118. 1701.07785s.

Skovgaard-Olsen, N., Singmann, H., Klauer, K.C., 2016. The relevance effect and conditionals. Cognition 150, 26–36. URL: http://dx.doi.org/10.1016/j.cognition.2015.12.017, doi:10.1016/j.cognition.2015.12.017.

Sloman, S., 2005. Causal Models How People Think about the World and Its Alternatives. Oxford University Press.

Sloman, S.A., Lagnado, D., 2015. Causality in Thought. Annual Review of Psychology 66, 223–247. URL: http://www.annualreviews.org/doi/abs/10.1146/annurev-psych-010814-015135, doi:10.1146/annurev-psych-010814-015135.

Spirtes, P., Glymour, C., 1991. An Algorithm for Fast Recovery of Sparse Causal Graphs. Social Science Computer Review 9, 62–72.

Takahashi, T., Kohno, Y., Oyo, K., 2010. Causal induction heuristics as proportion of assumed-to-be rare Instances (pARIs), in: Proceedings of the 7th International Conference on Cognitive Science (ICCS2010), pp. 361–362.

Takahashi, T., Oyo, K., 2014. Biconditional probability as a model of symmetry inference and causal induction from small samples, in: Proceedings of the 31st Annual Meeting of the Japanese Cognitive Society, pp. 141–148.

Tanaka, K., Namiki, N., Oyo, K., Takahashi, T., 2015. Causal induction by observation with uncertain occurrences, in: Proceedings of the 29th Annual Conference of the Japanese Society for Artificial Intelligence, pp. 1E3–1in.

White, P.A., 2003. Making Causal Judgments From the Proportion of Confirming Instances: The pCI Rule. Journal of Experimental Psychology: Learning, Memory, and Cognition 29, 710 –727.

Yokokawa, J., Takahashi, T., 2014. Inductive inference of causal relationship from uncertain co-occurrence information, in: Proceedings of the 76th National Convention of IPSJ, pp. 1T–6.

